# On inputs to deep learning for RNA 3D structure prediction

**DOI:** 10.1101/2025.02.14.638364

**Authors:** Marcell Szikszai, Marcin Magnus, Sachin Kadyan, Elena Rivas

## Abstract

Today, there are several effective deep learning models for predicting the 3D structure of proteins. Building on their success, models have been developed for predicting the 3D structure of non-coding RNAs. Unfortunately, these models are much less accurate than their protein counterparts. In this paper, we highlight differences between protein and RNA structure, and demonstrate methods for deep learning targeted at addressing those differences, with the aim of prompting discussion on these topics. We present an RNA-specific pipeline for generating structural Multiple Sequence Alignments (MSAs). Derived from the structural alignments, we introduce engineered evolutionary features that strongly inform RNA structure. Further, from the crystal structure, we derive structural features describing RNA base pairing. These evolutionary and structural features can be used in loss functions at different stages of training. Finally, we discuss different cropping strategies informed by RNA structure.

## 1 Introduction

The prediction of 3D protein structure was revolutionized via deep learning by AlphaFold in 2018 (Senior et al., 2020). Since then, the number of tools that apply deep learning to broader problems in structural biology has skyrocketed. It is no surprise that researchers immediately began adapting the lessons from AlphaFold toward non-coding ribonucleic acid (RNA) 3D structure prediction (Chen et al., 2020; Wang et al., 2021; Sato et al., 2021; Fu et al., 2021; Pearce et al., 2022; Shen et al., 2022; Baek et al., 2022; Feng et al., 2022; Li et al., 2022; Abramson et al., 2024). The core problems are essentially analogous: take a 1D polymer sequence as input, and predict the 3D conformation of the molecule. For both proteins and structured RNAs, the 3D structure is a consequence of the 1D polymer sequence, and the 3D structure has strong ties to the molecule’s function.

With increasing interest in 3D prediction of the RNA structure, there was a need for more robust tools to benchmark performance. The blind-assesment competition CASP15 (Kryshtafovych et al., 2023) joined RNA-Puzzles (Cruz et al., 2012; Miao et al., 2015; 2017; 2020) in 2022 to include RNA-only targets in the competition. These assessments of novel RNA structures indicates that to this day, deep-learning methods have yet to catch up to other existing traditional methods for RNA structure prediction, as reported by the latest CASP16 and RNA-Puzzle (Miao et al., 2020) competitions.

The amount of RNA structural data available to perform deep-learning RNA 3D structure prediction pales in comparison to that available for protein structure prediction (Szikszai et al., 2024). Methods like RNA3DB (Szikszai et al., 2024) have been created recently to exhaustively characterize structural homologies in existing RNA PDB chains, and to provide flexible tools to avoid structural homology overlap when designing training and testing sets for robust benchmarking.

As we learn from the successes in protein structure prediction, several additional considerations have to be taken into account when creating deep-learning methods for RNA structure prediction, owing to the distinct properties of RNA compared to proteins. We investigate some of these considerations in detail in this manuscript. Our contribution is a method to generate RNA structural alignments, and evolutionary and structural features that can be used to inform the training of RNA 3D structure prediction methods. See Figure 1.

**Figure 1:**
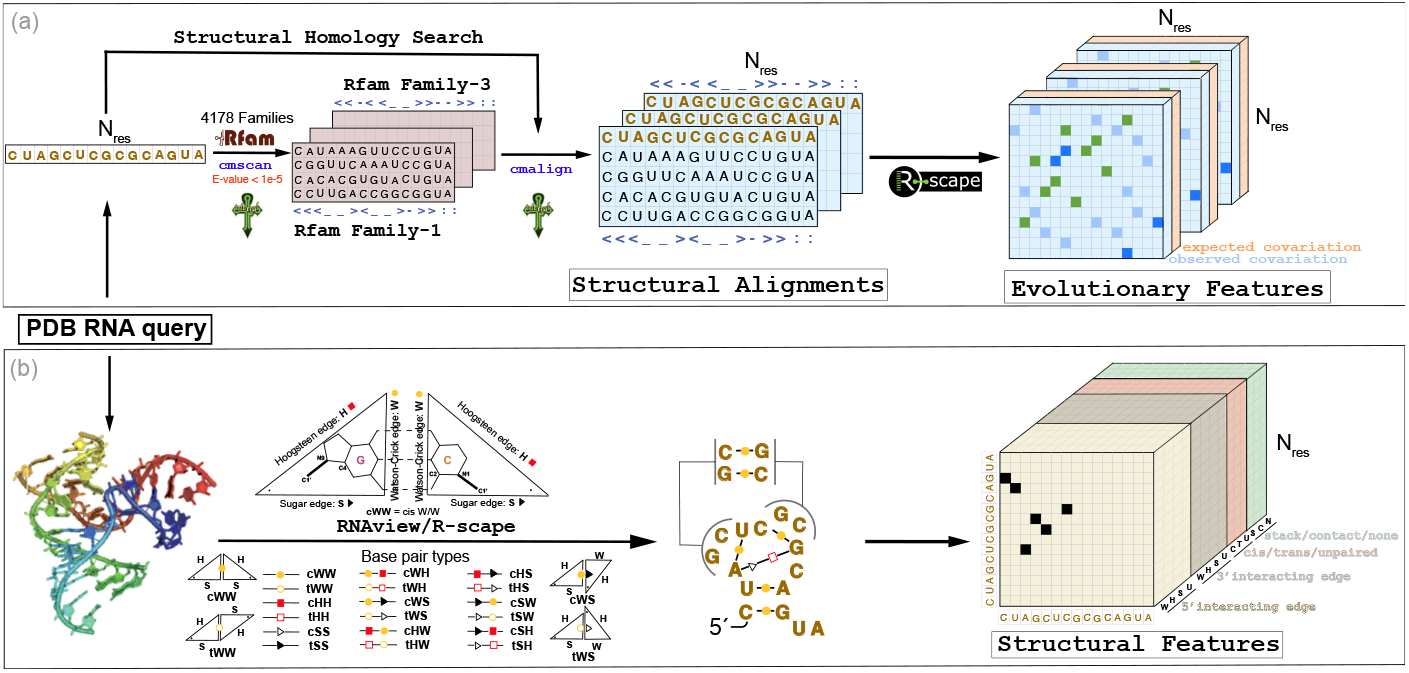
Overview of RNA structural inputs for training deep-learning methods. **(a)** The structural alignments are generated by structural homology search to the Rfam database (Ontiveros-Palacios et al., 2024), with the Infernal (Nawrocki & Eddy, 2013) method. We build one structural alignment for each Rfam family with significant homology to the query (E-value < 1e-5). See Appendix section B for details. From each structural alignment, we extract evolutionary features (both observed and expected covariation) using R-scape (Rivas et al., 2017; 2020). **(b)** From the query PDB file, we extract structural properties related to the base pairing geometry (including all possible base-pair types) using RNAview (Yang et al., 2003).

## 2 Rfam-based structural multiple sequence alignments

The idea behind using multiple sequence alignments (MSAs) as the input to AlphaFold-like deep learning models is the expectation that evolutionary information informs structure. This is true for both proteins and RNAs. However, with RNAs, the degree to which alignments inform structure varies highly by type. Structured RNAs, such as tRNA and rRNA, rely on specific conformations to perform their function. As a result, their structure is highly conserved. This conservation is easily detectable in their MSAs. Base pairing in particular can be inferred from alignments by looking at positive and negative evolutionary information (Rivas, 2020). On the other hand, some RNAs, like mRNAs, rely primarily on their conserved codon organization to determine their function. These mRNAs still form base pairs and fold into some 3D conformation, but they are generally not conserved (even though certain folded configurations may be more stable).

Since most interest in RNA 3D structure is focused on structured RNAs, alignments should be made with evolutionary conservation of structure in mind. However, the methods used by current deep learning models either assume all RNAs are structural, or do not incorporate structural conservation into their alignments. There are, broadly speaking, two pipelines representative of the approaches used by all existing methods: the rMSA (Zhang et al., 2023) pipeline that fits alignments to proposed structures, or structure-agnostic HMMER-based (Eddy, 2008; 2009; 2011) pipelines as used by AlphaFold 3.

The pipeline used by AlphaFold 3 relies on a large database of clustered representative RNA sequences from Rfam (Ontiveros-Palacios et al., 2024), RNAcentral (RNAcentral Consortium, 2021), and Nucleotide Collection (Sayers et al., 2023). HMMER is then used to find homologous sequences in this database. HMMER uses profile hidden Markov models to search sequence databases for homologues using sequence only. It does not consider the secondary structure of RNAs in the homology search.

It is well established that alignments for structural RNAs can be improved by using both sequence and secondary structure (Freyhult et al., 2007; Nawrocki, 2009), using structural homology methods such as Infernal (Nawrocki & Eddy, 2013). Infernal works by constructing profile stochastic context-free models (*covariance models* or CMs) of RNA families, which are trained from a family-specific MSA along with a consensus secondary structure.

The rMSA pipeline, used by RoseTTAFoldNA (Baek et al., 2023) and others (Wang et al., 2023), starts out with an initial HMMER alignment to first identify homologous sequences. Then it creates an Infernal covariance model using a predicted RNA secondary structure. This Infernal covariance model is then used for homology searches to arrive at a final alignment. This pipeline does consider secondary structure, but a relatively unreliable one, since the consensus is found through thermodynamic folding (Lorenz et al., 2016). Importantly, this approach assumes that the query sequence conserves its secondary structure, which may not be the case (e.g. mRNAs or synthetic constructs). Additionally, Infernal is used with an E-value cut-off of 10, which is prone to favor the inclusion of false positive homologues. The artifacts created by the high E-value cut-off and the assumption of conserved secondary structure was previously documented by Gao et al. (2022).

For known structural RNAs used in training, we propose to take advantage of the Rfam database (Ontiveros-Palacios et al., 2024). Rfam compiles a database of Infernal covariance models which classifies structural RNAs into families. Each family has a *seed alignment*, the MSA used to build the covariance model, along with a carefully created consensus secondary structure. Our method described in Figure 1a starts by finding Rfam families that show statistical significant homology to the query RNA sequence. For each homologous RNA family, a structural alignment including the query and sequences that belong to the Rfam family can be constructed. For the PDB database, as reported by RNA3DB (2024-12-04-full-release)(Szikszai et al., 2024), of the 1,869 RNA representative chains (clustered at 99% identity), 67% (1,198) have homology to at least one Rfam family with an E-value cutoff of 1*e*^*−*5^. RNA chains without homology to Rfam usually fall in the category of synthetic sequences, mRNAs or small fragments lacking structure.

From our structural alignments, we can directly feed covariation evolutionary information into the model in order to provide a signal about secondary and tertiary structure. See Figure 1a and Appendix Section A for a discussion.

Figure 2 presents a comparison of the performance of the alignment methods on a 5S rRNA structural RNA chain and a Purine riboswitch aptamer. Even though the AlphaFold-like alignment includes many more sequences, the alignment is less accurate identifying the positions that are base paired. Similar results are presented in the supplement for two other structural RNAs. Other examples are given in Appendix Section B.

**Figure 2:**
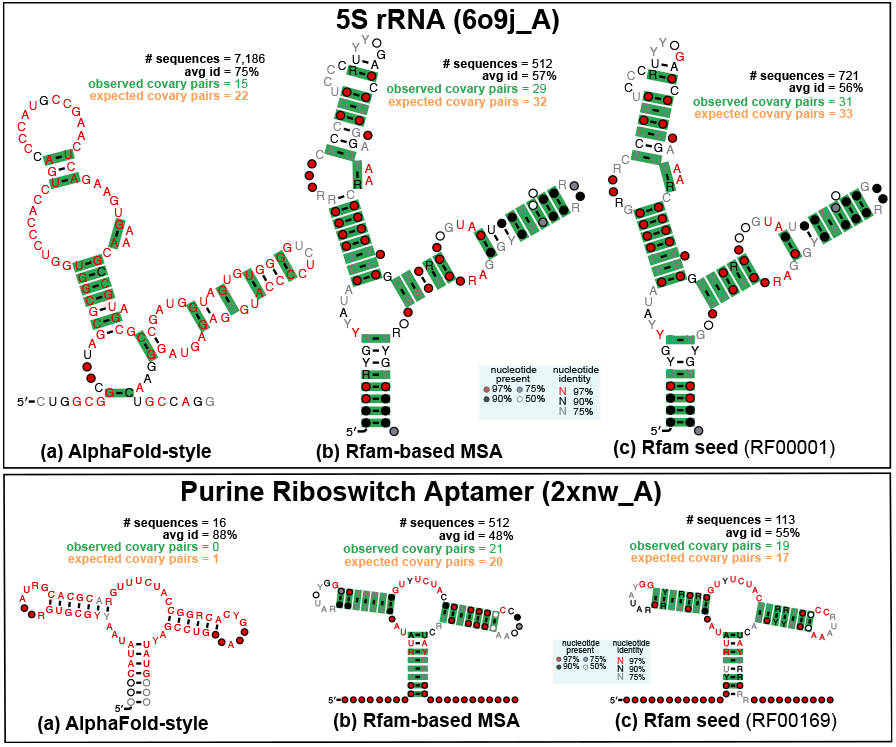
Comparison of alignment methods. For a 5S rRNA PDB chain and a Purine riboswitch aptamer, we show: **(a)** A HMMER alignment made against Rfam, RNAcentral and the nucleotide databases. **(b)** A structural alignment created with our Rfam-based method. **(c)** A curated structural alignment for the RNA family from Rfam. We show the evolutionary information present in each alignment as the significantly covarying base pairs (depicted in green) found using R-scape. Covarying base pairs are given in the context of a CaCoFold (Rivas, 2020) consensus secondary structure that incorporates all significantly covarying pairs found in the alignment.

These examples show the power of using Rfam’s covariance models, and building MSAs using Infernal with appropriate E-value cutoffs. In cases where no Rfam CM produces a significant hit, it may be difficult to determine whether the RNA is structural or not. As a result, alignments made with HMMER (as done by AlphaFold 3) will be the most adequate and informative without making assumptions about a secondary structure that could introduce circular analysis artifacts (Gao et al., 2022).

## 3 Loss functions associated to RNA base pairing

RNA folding is hierarchical (Tinoco & Bustamante, 1999) meaning that the secondary structure (that is, the collection of base pairs) is more stable and forms faster than the 3D structure (Banerjee et al., 1993; Mathews et al., 1997; Onoa et al., 2003). The RNA secondary structure heavily informs the 3D (Shapiro et al., 2007; Miao et al., 2020). As a result, it is often argued that for a model to predict 3D correctly, it must correctly predict the 2D structure first (Kerpedjiev et al., 2015). However, there is little discussion from the RNA 3D structure prediction community about loss functions that target RNA base pairing specifically.

AlphaFold-like models distinguish two different kinds of losses. Structural losses rely on the entire 3D structure. These are usually end-to-end, and are evaluated at the end of the structure module or the diffusion module in the case of AlphaFold 3, e.g. Frame Aligned Point Error (FAPE) (Jumper et al., 2021). Also, there are auxiliary losses that apply to linear projections from the internal pair representation, usually just before the structure module and after the Pairformer or Evoformer, e.g. distogram loss (Jumper et al., 2021). Here we propose two loss functions, one structural and one auxiliary to inform specifically about RNA base pairing.

We propose a loss termed *pairtogram* loss–a play on AlphaFold’s distograms^1^. A pairtogram describes the base pairing geometry and can be used as an auxiliary loss. To construct a pairtogram, we extract an augmented Leontis-Westhof base pair geometry classification matrix (Leontis & Westhof, 2001) for all *N*_res_ *× N*_res_ pairs. This classification provides 12 basic geometric types, distinguishing between Watson-Crick, Hoogsteen, or Sugar-edge interacting edges, and cis or trans bond orientations. For instance, canonical RNA base pairs `A:U`, `G:C` are cis interactions between the Watson-Crick edges of both residues. The standard annotation is augmented with whether the pair is stacked (but not any of the defined base pairs), or a contact (defined as residues at a distance smaller than 8 Å), or neither. See Appendix section C for details on how the pairtogram loss is calculated.

Our RNA structural loss considers the dihedral angle between the planes of the two nucleotide bases. Pyrimidine bases are completely planar, while the base in purines is nearly planar (but can be functionally treated as such) (Callahan, 2011). As a result, we can assign a plane corresponding to a base via three atoms in all residues. In principle, any three atoms can be used since the bases are largely planar, but we consider two planes for both pyrimidines and purines for redundancy. The planes for purines are defined by C1^*′*^-N9-C4 and C8-N9-C4, while the planes for pyrimidines are defined by C1^*′*^-N1-C2 and C6-N1-C2 (see Figure 3). These planes are then used to calculate the dihedral angles between bases. Residues forming Watson-Crick base pairs will have angles close to 180^*°*^. Moreover, residues in one side of a canonical helix will also have angles close to 180^*°*^ with residues on the other side of the helix. On the other hand, residues in one side of the helix will have very small base angles amongst each other.

**Figure 3:**
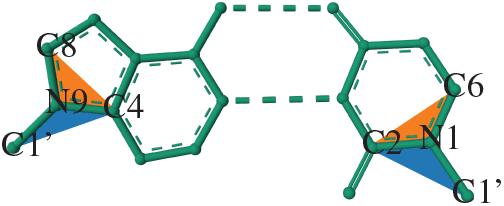
Between-Base Angle Planes. An example of how the planes are constructed for a purine (left, adenine) and a pyrimidine (right, uracil). The first planes are shown in blue, while the second planes are shown in orange. The example base pair is from a tRNA (PDB: `1ehz_A`) (Shi & Moore, 2000) and drawn with Mol* (Sehnal et al., 2021).

Let the plane for a residue *i* be defined by three atom coordinates (**A**_C1’/C8/C6_, **A**_N9/N1_, **A**_C4/C2_) where **A** *∈* ℝ^3^. Then, we calculate the normal of the plane,

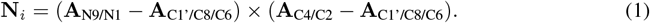

For each pair of planes *i* and *j*, we calculate the sine and cosine of the dihedral angle,

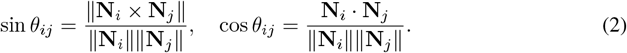

Then we define our loss, Between Base Angle Error (BBAE) as,

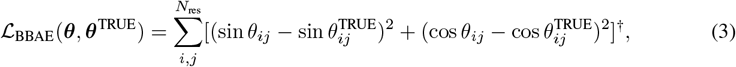

where ***θ*** are the dihedral angles for the predicted structure, and ***θ***^TRUE^ are the dihedral angles for the ground-truth structure.

## 4 Residue cropping

A key consideration for deep learning models is memory. Recently, this issue has received a lot of attention, since large-scale state-of-the-art models (such as large language models) may have hundreds of billions of parameters (Kaplan et al., 2020; Grattafiori et al., 2024), which must be loaded into memory during both inference and training. In the case of 3D structure prediction models, the number of parameters is much more manageable (on the order of around 100M parameters for AlphaFold 2 (Jumper et al., 2021), for example), but the memory consumption is still relatively high, and can scale cubically or quadratically with sequence length (Vaswani et al., 2017; Senior et al., 2020; Jumper et al., 2021). As a result, deep learning models for 3D structure prediction often have to crop the sequences to shorter sub-sequences during training and process these crops one at a time. This is commonly referred to as *residue cropping*.

Initially, the AlphaFold 1 convolutional neural network (CNN) produced crops of length 64 in order to generate 64 *×* 64 distograms (Senior et al., 2020). This cropping strategy is easy to motivate for proteins: existing literature has shown that protein contact prediction only needs a limited context window (Jones & Kandathil, 2018; Senior et al., 2020). Unfortunately, this is not the case for RNAs. Some structural RNA families, such as SSU and LSU rRNA subunits, contain long-range base pairs that are more than 500 residues apart, and are difficult to predict for traditional dynamic programming algorithms (Huang et al., 2019).

In 2021, AlphaFold 2 moved from a CNN-based architecture to a transformer model, which requires *O*(*n*^3^) memory for a sequence of length *n*. This memory requirement comes from *triangular* selfattention. As a result, AlphaFold 2 takes highly restrictive crops of 256 residues during the initial training phase and crops of 384 residues during fine-tuning. The starting position of these crops is randomly sampled from *𝒰* {1, *N*_res_ *−* `crop_size` + 1*}* where *N*_res_ is the length of the sequence.

These crops are *contiguous* in sequence, so that given a starting position *i*, the window contains residues [*i, i* + `crop_size` *−* 1].

While this strategy works well for proteins, since they only need small context windows, it is suboptimal at best for RNAs. AlphaFold 2 contiguous crops break RNA base pairs, and include only one half of the RNA canonical helices that are the foundation of any RNA 3D structure (Figure 4a). Despite this, some methods for RNA 3D structure prediction, such as DRfold (Li et al., 2022) use continuous cropping for training. Consequently, DRfold also restricts their test set to RNAs with lengths between 14 and 392 nucleotides (Li et al., 2022), likely masking the performance degradation of their cropping strategy.

**Figure 4:**
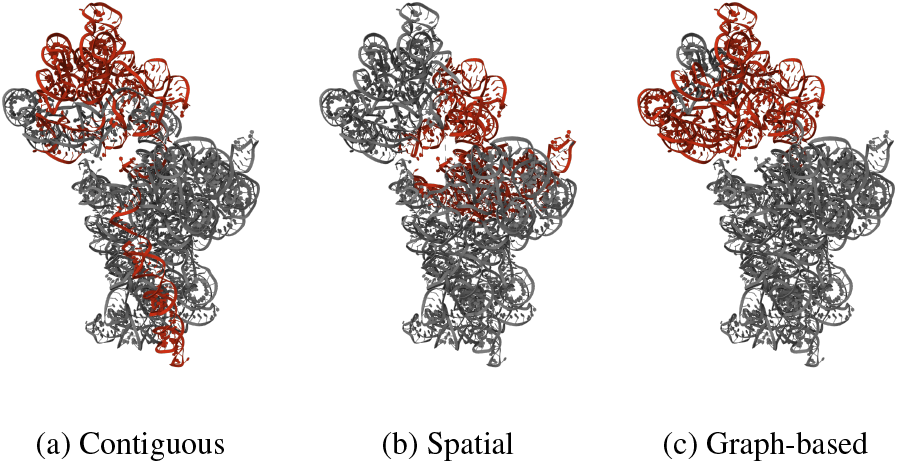
Different cropping strategies. Example for a 30S rRNA (PDB: `5no2_A`) (López-Alonso et al., 2017). The red regions indicate the cropped sequence. The starting position is 1,100 with `crop_size` = 384, with Watson-Crick base pairs extracted using RNApdbee 2.0 (Zok et al., 2018) and RNAview (Leontis & Westhof, 2001). The visualisations were created with Mol* (Sehnal et al., 2021).

When DeepMind debuted AlphaFold-Multimer for protein complex prediction in 2022, they introduced *spatial* crops where residues are selected by their spatial distance in the 3D structure. For these spatial crops, a starting position is sampled from 𝒰 {1, *N*_res_}. A reference atom is chosen (in the case of AlphaFold-Multimer C_*α*_ atoms), from which all distances are measured. Then, the `crop_size` nearest residues, measured by Euclidean distance to the reference atoms, are taken as the crop (Figure 4b). In AlphaFold-Multimer, spatial crops are chosen in a 50:50 ratio along with contiguous crops with a `crop_size` of 384. Although only used by AlphaFold-Multimer for multimer interface residues, the concept can be easily adapted to monomers.

Beyond *spatial* crops, RoseTTAFoldNA (Baek et al., 2023) also developed an alternate strategy for cropping nucleic acid–protein complexes and RNA monomers to explicitly avoid breaking base pairs and to pick context windows better suited to preserve RNA canonical helices composed of stacked Watson-Crick base pairs (Figure 4c). For RNA monomers, RoseTTAFoldNA builds a weighted undirected graph of the sequence where sequential residues have edges with a weight of one, and Watson–Crick base pairs have a weight of zero. As before, a random starting position is sampled from 𝒰 *{*1, *N*_res_ *}*, and minimum-weight graph traversal is used to find the nearest `crop_size` = 256 residues based on the graph-distance.

We suggest a combined method using *contiguous* cropping together with RoseTTAFoldNA’s *interaction graph-based* and AlphaFold 3 *spatial* crops. The ratios of the different cropping categories can be customized for the different training sets and different environments, with the goal of balancing time and performance with structure prediction accuracy.

## 5 Summary

The prediction of RNA 3D structure from sequence by deep learning methods is challenged by the small amount of structural data existing to train the models in comparison to proteins. This manuscript aims at lowering the impact of such a fundamental problem by making sure that the information obtained from the existing inputs extracts the maximal amount of structural RNA specific properties, both structural and at the level of RNA base pairing.

We have presented structural alignments for PDB RNA chains that capture significantly more pairing information than other agnostic homology methods. We have introduced losses capturing RNA base pairing information, including non-canonical base pairs, and also structural losses that capture the stacked nature of the 3D helices formed by the RNA canonical base pairs. Finally, we have discussed base pairing-aware cropping strategies. By introducing these topics in this manuscript, we hope to encourage further research and discussion on RNA-specific models in the field of RNA 3D structure prediction.

## A Structural Evolutionary features

As discussed in Section 2, the purpose of feeding MSAs into deep learning structure prediction pipelines is to provide evolutionary context about the residues. In traditional RNA structure prediction pipelines, MSAs can allow the model to identify covariation resulting from the presence of conserved RNA structure. For structured RNAs, the covariation derived from their alignments has been shown to be highly informative towards both the secondary and tertiary structures (Rivas et al., 2017; Rivas, 2020; Rivas et al., 2020; Rivas, 2023; Karan & Rivas, 2024).

Here we describe a method for how these features can be used by an AlphaFold-like architecture. Using the software R-scape (Rivas et al., 2017; 2020; Rivas, 2020), for any pair of positions, we can compute both the statistically significant covariation above phylogenetic expectation or observed covariation (in the form of an expected E-value), as well as the expected covariation (or power) given the number of total substitutions in the pair. Both E-value and power are binned into 8 bins, and E-value is clamped to the range [0, 10.0]. The feature tensor produced is of shape [*N*_res_, *N*_res_, 16]. This is then linearly projected to the pair representation channel dimension *c*_*z*_, and added to the input of, for example, the main transformer block. This method also allows for calculating the E-value and power from multiple MSAs by computing the minimum, mean, and maximum across the MSAs, and producing a tensor of shape [*N*_res_, *N*_res_, 16 *×* 3].

Under ideal circumstances, it may seem unnecessary to explicitly feed in these features. We may expect that since deep learning is highly effective at representation learning, i.e. the ability to learn the useful representations from the raw data, we can just directly input the raw alignments and the network can learn these features implicitly as part of its internal representation. Currently we have no evidence to conclusively show whether inputting these features directly improves performance, however, there are two other possible motivations for explicitly calculating these.

First, these evolutionary maps provide an efficient way of embedding covariation information from a high-dimensional MSA in a way that is independent of the MSA dimensionality. AlphaFold 2’s Evoformer memory cost is 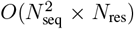, where *N*_seq_ is the number of sequences in the alignment^3^. To address this, the model reduced the depth of the alignments using *MSA clustering*, where a relatively small number of sequences (*N*_clust_ = 128 during the initial training phase, *N*_clust_ = 512 during fine-tuning^4^) cluster centres are picked. Then the remaining sequences in the MSA are assigned to their closest cluster by Hamming distance, and a number of statistics (e.g. distribution of amino acids) are computed for the cluster. Our proposed evolutionary features avoid the effect of any dimensionality reduction.

Second, while we currently suggest using these features as inputs, they may also be useful as auxiliary losses. It would be easy to create an auxiliary head that linearly projects the internal representation into the desired dimension (either [*N*_res_, *N*_res_, 16] or [*N*_res_, *N*_res_, 16 *×* 3]), and calculates the averaged cross-entropy loss as done for distograms by AlphaFold 2. We further suggest that these features may also be a useful auxiliary head for other models such as RNA language models like RiNALMo (Penić et al., 2024).

## B Structural alignments

For our Rfam-based MSAs, each family-specific alignment includes the query sequence, and up to 256 seed sequences from the Rfam family, or up to 512 full sequences if there are fewer than 256 seed sequences in the family.

In Figure 5, we describe two other examples of structural RNAs and the comparison of our Rfam/Infernal based structural alignments and the structure-agnostic alignments used by AlphaFold 3. For the two selected PDB chains: `5ddp_A` (a glutamine riboswitch aptamer) (Ren et al., 2015) and `2qus_A` (a Hammerhead_3 ribozyme) (Chi et al., 2008), we observe that Infernal is able to find multiple homologs in the Rfam database, and produces a structural alignment of quality comparable to that of the Rfam seed. On the other hand, a structure-agnostic search in the same database renders few homologs and very sequence-conserved alignments that offer no evolutionary information about the secondary structures.

**Figure 5:**
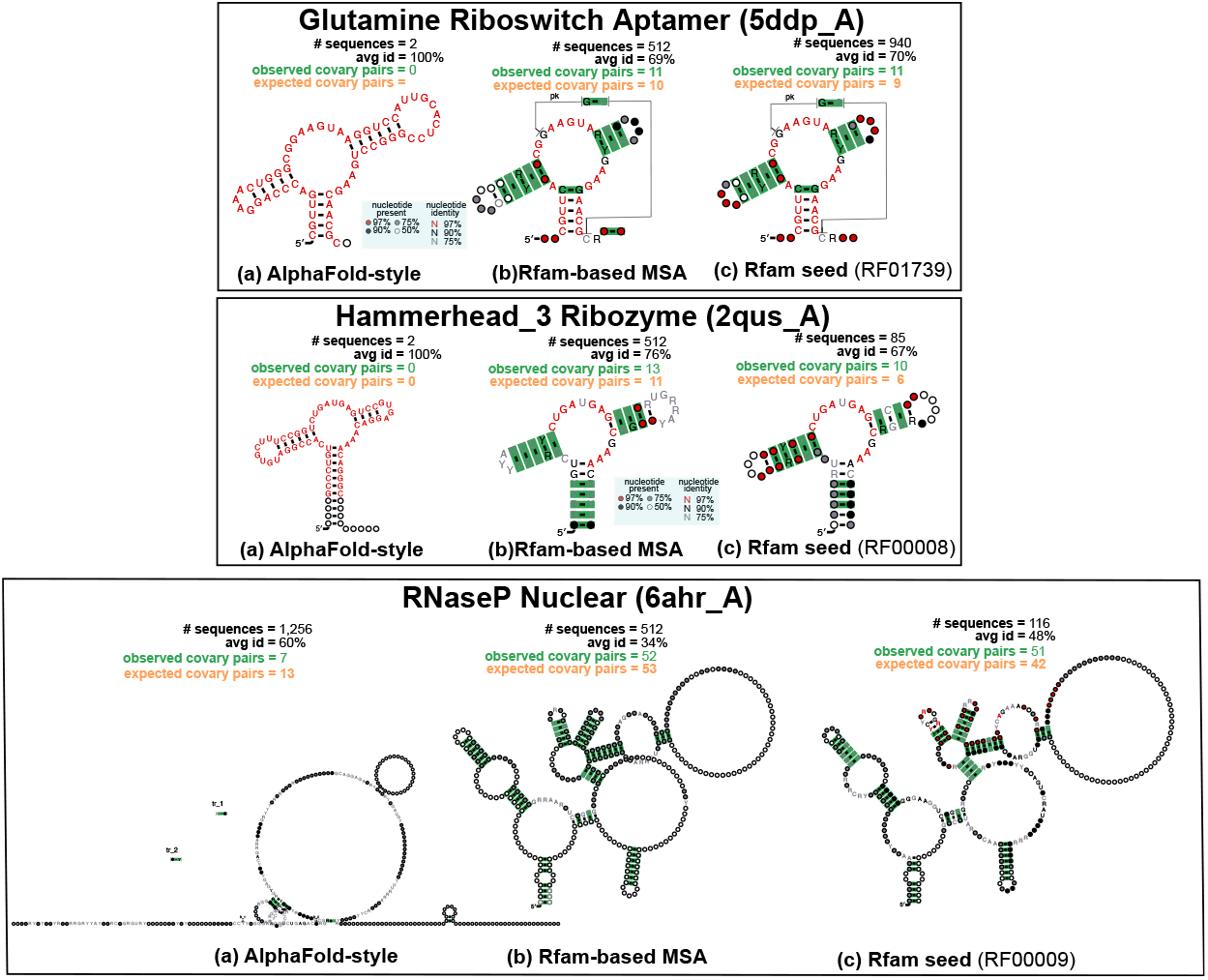
Comparison of alignment methods. **(a)** The AlphaFold 3-like alignments were created in-house using the following database: Nucleotide collection (nt) 112,177,963 sequences; 2,688,129,930,104 total bases (Feb 2, 2025 4:42 AM); BLASTDB Version: 5; RNAcentral Release 24, 07/03/2024. **(b)** Our structural alignments were created using Rfam v15.0, and Infernal v 1.1.4. **(c)** We compare to the Rfam seed alignment for the RNA family to which the queries belong. Evolutionary conserved base pairs are depicted in green. For other details, see caption of Figure 2.

## C Pairtogram loss details

Pairtogram data is extracted from the 3D structures using R-scape (Rivas et al., 2017; 2020; Rivas, 2020), which includes a modified version of the software RNAView (Leontis & Westhof, 2001). The final pairtogram matrix has 14 total dimensions for each pair (see Table 1 and Figure 1b for a breakdown of the features). We treat these 14 dimensions as four separate one-hot vectors, and calculate the average cross-entropy loss across the four features between the ground-truth and a linear projection from the pair representation.

**Table 1:** Structural features used in the *pairtogram* loss. The final tensor has size [*N*_res_, *N*_res_, 14].

| Description | Values | Dimension |
| --- | --- | --- |
| 5' interacting edge | Watson-Crick, Hoogsteen, Sugar-Edge, unpaired | 4 |
| 3' interacting edge |  | 4 |
| Bond orientation | cis, trans, unpaired | 3 |
| Stacked/contact | stacked, contact, neither | 3 |

## Footnotes

1 *Distance histograms*, an output of discretised pairwise distances between all atoms.

3 This is a simplification from the original AlphaFold 2 paper. We omit templates from our explanations.

4 This is a simplification. *N*_seq_ = *N*_clust_ + *N*_templ_.

† Note that 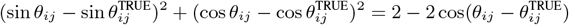.

