## Supplementary figures and images for "On inputs to deep learning for RNA 3D structure prediction"

### 2qus_A_1.cacofold.R2R.sto.pdf

#### 2qus_A_1.cacofold

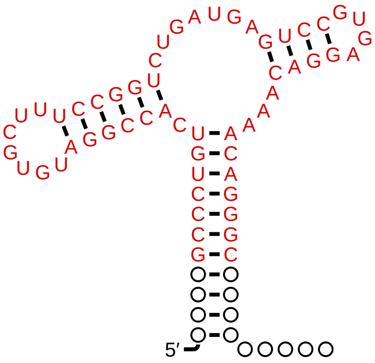

### 2xnw_A_1.cacofold.R2R.sto.pdf

#### 2xnw_A_1.cacofold

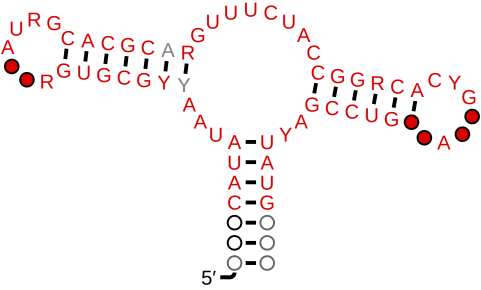

### 5ddp_A_1.cacofold.R2R.sto.pdf

#### 5ddp_A_1.cacofold

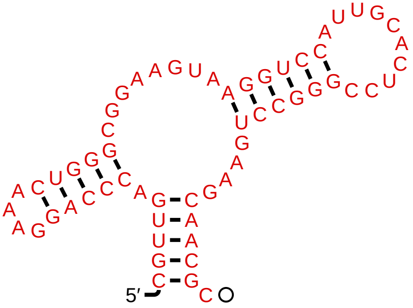

### RF00008_4_59_1.cacofold.R2R.sto.pdf

#### RF00008_4_59_1.cacofold

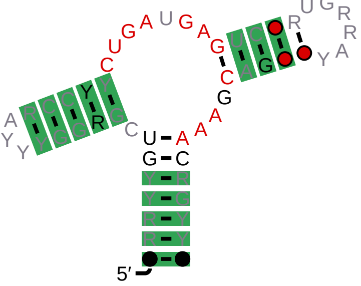

### RF00008_4_59_1.R2R.sto.pdf

# RF00008_4_59_1

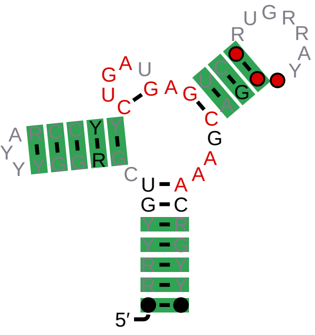

### RF00008_Hammerhead_3.cacofold.R2R.sto.pdf

#### RF00008_Hammerhead_3.cacofold

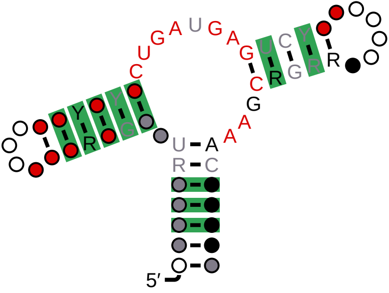

### RF00008_Hammerhead_3.R2R.sto.pdf

#### RF00008_Hammerhead_3

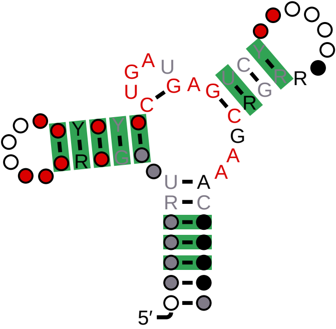

### RF00167_0_66_1.cacofold.R2R.sto.pdf

#### RF00167_0_66_1.cacofold

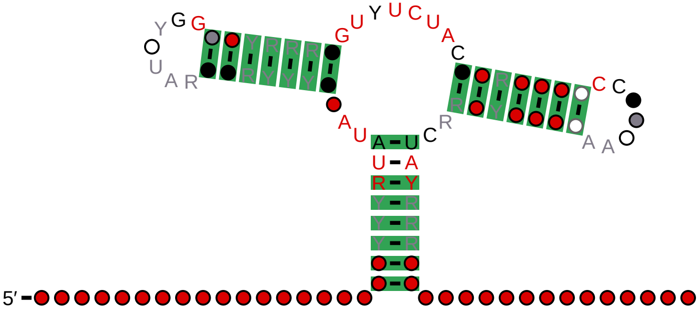

### RF00167_0_66_1.R2R.sto.pdf

# RF00167_0_66_1

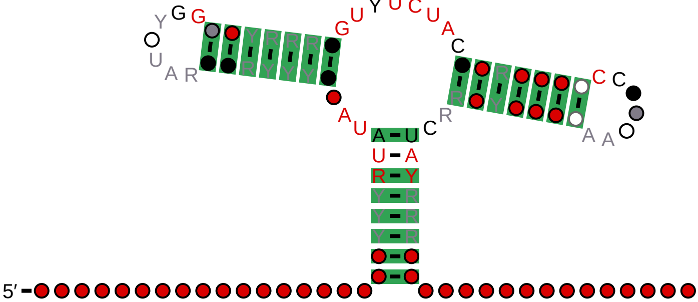

### RF00167_Purine.cacofold.R2R.sto.pdf

#### RF00167_Purine.cacofold

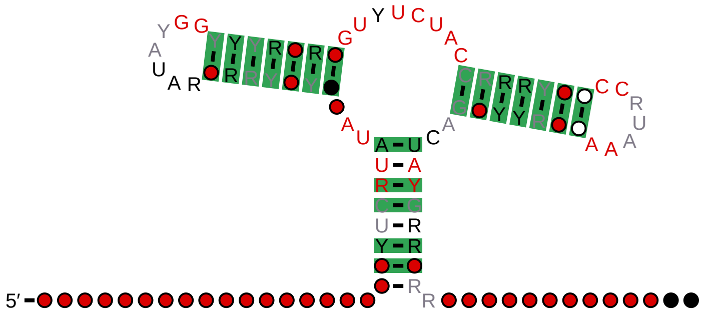

### RF00167_Purine.R2R.sto.pdf

#### RF00167_Purine

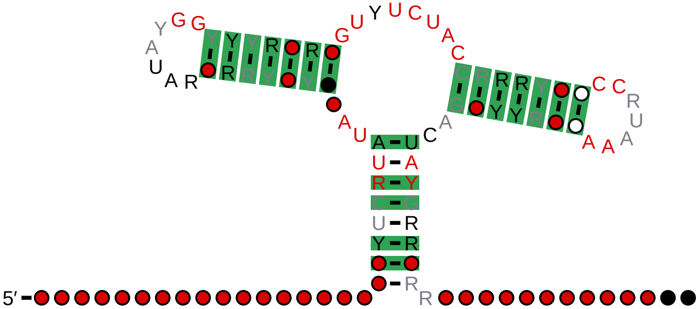

### RF01739_0_60_1.cacofold.R2R.sto.pdf

#### RF01739_0_60_1.cacofold

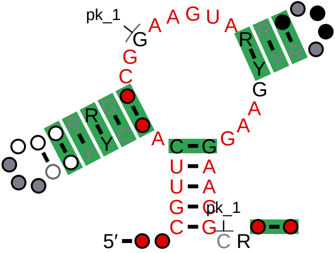


# pk_1

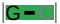

### RF01739_0_60_1.R2R.sto.pdf

# RF01739_0_60_1

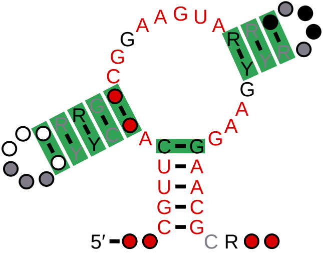

### RF01739_Glutamine.cacofold.R2R.sto.pdf

#### RF01739_Glutamine.cacofold

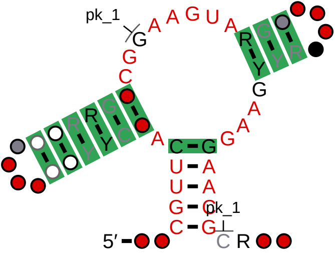


# pk_1

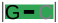

### RF01739_Glutamine.R2R.sto.pdf

#### RF01739_Glutamine

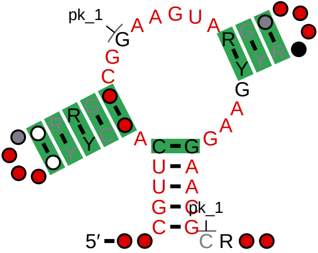


# pk_1

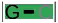
